# Genetic modification of *Clostridium kluyveri* for heterologous *n*-butanol and *n*-hexanol production

**DOI:** 10.1101/2025.05.05.652178

**Authors:** Caroline Schlaiß, Saskia T. Baur, James W. Marsh, Kurt Gemeinhardt, Largus T. Angenent

## Abstract

The mesophilic microbe *Clostridium kluyveri* serves as the most commonly used model microbe to elucidate the physiology and biochemistry of ethanol-based chain elongation *via* reverse ß-oxidation. In this pathway, ethanol and acetate are converted into short- and medium-chain carboxylates. However, to date, no genetic system has been published in a peer-reviewed publication. Here, we report the development of versatile genetic tools for *C. kluyveri*, utilizing the pMTL *Clostridia* shuttle vector system and thiamphenicol as a selective marker. We identified the native restriction-modification system of *C. kluyveri* as a critical barrier to DNA transfer and overcame it by identifying and characterizing the crucial methyltransferase. To mimic the native DNA methylation pattern of *C. kluyveri*, we performed *in-vivo* methylation of the shuttle vector plasmid by expressing the methyltransferase in *Escherichia coli*, followed by DNA transfer *via* conjugation. After validating the genetic system, we demonstrated heterologous expression of different combinations of both NADH and NADPH-dependent alcohol dehydrogenases from *Clostridium acetobutylicum*. The expression of these genes was controlled by the P*_thl_* promoter, which is commonly used in *Clostridia,* and the P*_adhE2_* promoter, leading to *n*-butanol and *n*-hexanol production of the mutant strains. This genetic system for *C. kluyveri* will not only enable further research on the metabolism of this microbe but also enable more profound insights into ethanol-based chain elongation in general.

**IMPORTANCE:** Medium-chain carboxylates are required in various everyday products, including cosmetics, pharmaceuticals, and fragrances, and show a natural antimicrobial property. Further, they represent food additives and serve as chemical building blocks for several other compounds. Traditionally, these carboxylates are produced from fossil resources, contributing to increased greenhouse gas emissions. Alternatively, they are derived from animal- or plant-based fat (*e.g*., coconut oil), which competes with agricultural land that is needed for food production. However, microbial chain elongation, which is a biotechnological approach relying on microbes, such as *Clostridium kluyveri*, is sustainable and a promising alternative to the conventional production of medium-chain carboxylates. Notably, it enables the use of industrial waste streams (*e.g.*, off-gases, carbohydrate-rich industrial waste) as substrates, making the process more environmentally friendly. By applying our genetic system for *C. kluyveri*, a better understanding of microbial chain elongation can be achieved, and potentially even an extension of its product portfolio.

## INTRODUCTION

Despite advances in renewable energy technologies, modern society remains heavily dependent on fossil fuels. This continued reliance contributes considerably to greenhouse gas emissions, which in turn accelerate climate change and the depletion of natural resources (1). Therefore, moving toward a more circular economy represents a critical challenge for sustainability to ensure long-term resilience for future generations. A key aspect of this transformation is the development of sustainable alternatives for essential chemical building blocks, which are conventionally assembled using fossil resources.

The development of sustainable alternatives represents a central challenge for modern environmental biotechnology. In particular, chemical building blocks that are in high demand from industry include short- and medium-chain carboxylates (SCCs and MCCs, respectively) (2). Both SCCs and MCCs contain a carboxylic group; while SCCs have carbon chains with a length of two to five carbon atoms, MCCs consist of six to twelve carbons (3) and their monetary value increases with an increasing chain length (4). MCCs serve as versatile chemical compounds with numerous industrial applications. Beyond functioning as essential building blocks for bioplastics (5, 6) and potential biofuel precursors through Kolbe electrolysis (7–9), these compounds have considerable utility across multiple sectors. They are widely incorporated into cosmetics, pharmaceuticals (2), fragrances (10), and rubber production (11). Additionally, MCCs feature prominently in food products as flavor additives (11, 12) and in animal feed formulations, where they provide valuable antimicrobial properties (13, 14). Also, they serve as precursors for alcohols such as *n*-butanol and *n*-hexanol (15).

Conventionally, MCCs are produced from petrochemicals (16), leading to environmental devastation and mobilization and release of carbon dioxide (CO_2_). A potential bio-alternative that is already applied on an industrial scale is the production of MCCs from animal or plant fat, for example, coconut oil, palm kernel oil, and castor oil (17, 18). However, the sustainability of this approach is contested because it requires agricultural land, which is indispensable for growing food, especially given the steadily increasing global population (19). Another promising biotechnological alternative is microbial chain elongation to produce MCCs.

By using the reverse ß-oxidation pathway, microbes fuse a C2 compound to short-chain carboxylates, resulting in a stepwise elongation of the carbon chain by two carbon atoms at a time, respectively, up to a maximum chain length of six carbon atoms (20). This elongation process can be achieved by reactor microbiomes using an open culture of microbial consortia. The open cultures do not require sterile treatment and are resilient to disturbances (21, 22). Another promising attempt, which was demonstrated in several studies, is to couple syngas fermentation with chain elongation by cultivating an acetogenic and a chain-elongating microbe in a co-culture (23–26). First, the acetogenic microbe produces the intermediates ethanol and acetate from a mixture of hydrogen, CO_2_, and carbon monoxide. Second, the chain-elongating microbe further elongates the intermediates to medium-chain carboxylates *via* reverse ß-oxidation. Third, the acetogenic microbe reduces SCCs and MCCs into their corresponding alcohol (*n*-butanol and *n*-hexanol). Besides open- and co-cultures, medium-chain carboxylate production performed by pure cultures offers higher production rates (27).

*Clostridium kluyveri* has been extensively used to study the physiology and biochemistry of ethanol-based chain elongation *via* the reverse ß-oxidation pathway (28–31). Therefore, it serves as a model microbe for microbial chain elongation. To date, no genetic system for this microbe has been published in a peer-reviewed study. Here, we developed a genetic system for *C. kluyveri* by mimicking its native methylation pattern on a shuttle vector plasmid to circumvent its restriction-modification defense system. As a proof of principle, we introduced and heterologously expressed alcohol dehydrogenases from *Clostridium acetobutylicum* to enable *n*-butanol and *n*-hexanol production. By applying the findings from this study, metabolic engineering of *C. kluyveri* can be used to increase its productivity, expand its product portfolio, and obtain more profound insights into the reverse ß-oxidation metabolism in general.

## RESULTS

### Plating optimization

To enable successful genetic modification, our initial focus was to optimize plating conditions for the reliable growth of single colonies, because a high plating efficiency increases the likelihood of isolating the correct clone. We systematically adjusted each variable to determine the optimal plating conditions. Of these, supplementing the DSMZ52 medium with a 10-fold concentration of vitamins and using cells in their early exponential growth phase had the most substantial positive impact on colony formation (**Table S1**). Other modifications, such as pour-plating with 0.8% Bacto agar, increasing the atmospheric pressure in the storage tanks to 0.6 bar, and placing a sterile filter paper (1 cm X 1 cm) soaked with ethanol in the lid of the Petri dish to ensure sufficient substrate supply, showed moderate positive effects (**Table S1**).

In contrast, using complex modified Reinforced Clostridial Medium (RCM) (32), which was supplemented with 20 mL/L ethanol and 2.5 g/L NaHCO_3_ instead of DSMZ52 medium, and adding ethanol to the external plate atmosphere affected the plating efficiency negatively (**Table S1**). Further, using paper clips to increase ventilation resulted in a drastic reduction in colony formation (**Table S1**), most likely due to ethanol evaporation and plate desiccation from the enhanced airflow. Because hydrogen production of *C. kluyveri* resulted in large bubbles of the pour-plated medium, which complicated post-treatment, we decided to perform spread-plating whenever a minor reduction in plating efficiency was acceptable. For routine plating, we chose DSMZ52 medium with increased vitamin supplementation over RCM to reduce contamination risk and improve consistency. However, RCM medium was necessary during conjugation to support the growth of *E. coli*.

### Antibiotic testing

Selection of cells carrying the desired plasmid requires two key factors: **1)** the sensitivity of the wild-type strain to the respective antibiotic, and **2)** the expression of an enzyme that ensures resistance to the antibiotic to which the wild-type strain is sensitive. Thiamphenicol resistance is a well-established marker in many clostridial strains, including strains closely related to *C. kluyveri* (33, 34). Therefore, we tested the sensitivity of the cells to thiamphenicol in liquid DSMZ52 medium and on plates and determined the minimum inhibitory concentration (MIC) after one week, which was 3 µg/mL (**Figure S1**). However, we chose to use a higher concentration of 5 µg/mL thiamphenicol in liquid medium to prevent the growth of wild-type cells due to spontaneous resistance or a reduction in antibiotic concentration caused by instability of the antibiotic at 37°C after prolonged incubation. Additionally, we used this concentration to select plasmid-carrying cells on solid media.

### Identification of the crucial methylation pattern

A restriction assay of the pMTL83151 shuttle vector plasmid (35), using the cell lysate of *C. kluyveri*, showed digestion of the foreign DNA, producing fragment sizes of approximately 2 kbp and 0.5 kbp (**Figure 1A**). This result indicated that the native restriction and modification defense system of *C. kluyveri* (**Figure 1B**) is a critical barrier that must be overcome for successful genetic modification. PacBio sequencing of the native methylation pattern of the genomic DNA of *C. kluyveri* DSM555 revealed methylation at the following DNA sites: GATC, GWTAAT, CCAAG, CAAAAAT, and CCGG, suggesting the presence of multiple restriction-modification systems. *In-silico* analysis revealed that the observed restriction pattern of the unmethylated pMTL83151 corresponded to the CCGG as the critical methylation site, corresponding with the findings of Schneider, 2024 (36).

**Figure 1.**
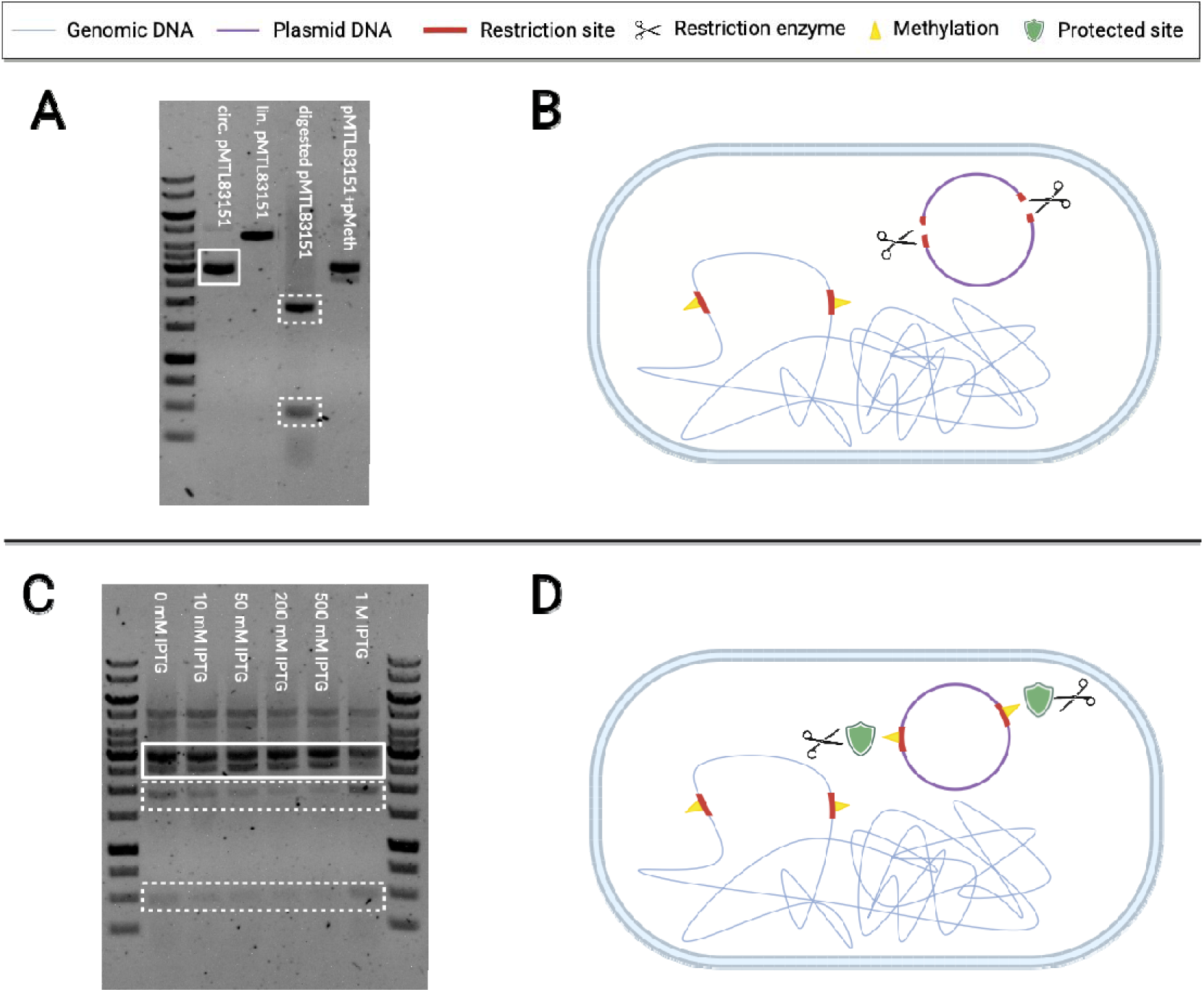
Restriction modification system of C. kluyveri. A) Restriction digest of the unmethylated pMTL83151 plasmid after 4 h. The solid-lined box in the 2^nd^ lane shows the unrestricted, circular plasmid (∼4.5 kbp), whereas the dashed-lined box in the 4^th^ lane shows the vector digested by C. kluyveri cell lysate (fragment sizes: ∼2 kbp, ∼0.5 kbp). The linearized pMTL83151 vector in the 3^rd^ lane (∼4.5 kbp) and the mixture of circular plasmids pMTL83151 and pMeth (5^th^ lane) serve as controls. B) In bacterial cells, DNA restriction serves as a defense mechanism against foreign DNA that either lacks methylation at the critical sites or shows a different methylation pattern from the native DNA of the respective bacterium. C) Restriction digest of C. kluyveri-specific methylated pMTL83151 after 24 h. Because the expression of the methyltransferase (CKL_2671) of C. kluyveri was controlled by the P_lac_ promoter, we tested different IPTG levels for induction (10°mM to 1 M). All induction levels result in minor restriction of pMTL83151 highlighted with dashed boxes (fragment sizes: ∼2 kbp and ∼0.5 kbp), however, the restriction was clearly reduced and minimized. Best results with the least restriction were obtained for the induction with 500 mM IPTG. D) As a result of the same methylation pattern as the native DNA, the foreign DNA is protected from restriction by the bacterial restriction modification system. Created with BioRender.com.

We identified a native methyltransferase in the REBASE database (37) that methylates CCGG sites and is encoded by the genome of *C. kluyveri* (locus tag CKL_2671). To counteract the restriction, we expressed this methyltransferase under the control of the Isopropyl-ß-D-thiogalactopyranoside (IPTG)-inducible P*_lac_* promoter on the plasmid pMeth (**Figure 2**) to mimic *C. kluyveri*-specific methylation of pMTL83151. As a result, the restriction assay testing CCGG-methylated pMTL83151 showed reduced plasmid digestion. Next, we optimized the methyltransferase expression by testing different IPTG concentrations (10-1000 mM) for induction in *E. coli* TOP10. In a final restriction digest (**Figure 1C**), we only observed minimal digestion of the shuttle vector after 24 h, indicating that the restriction sites of the foreign DNA were protected by methylation (**Figure 1D**). The lowest restriction activity occurred at IPTG induction concentrations between 200 mM and 500 mM right after inoculation. Based on these results, we established 500 mM IPTG as the standard concentration for inducing the expression of the methyltransferase.

**Figure 2.**
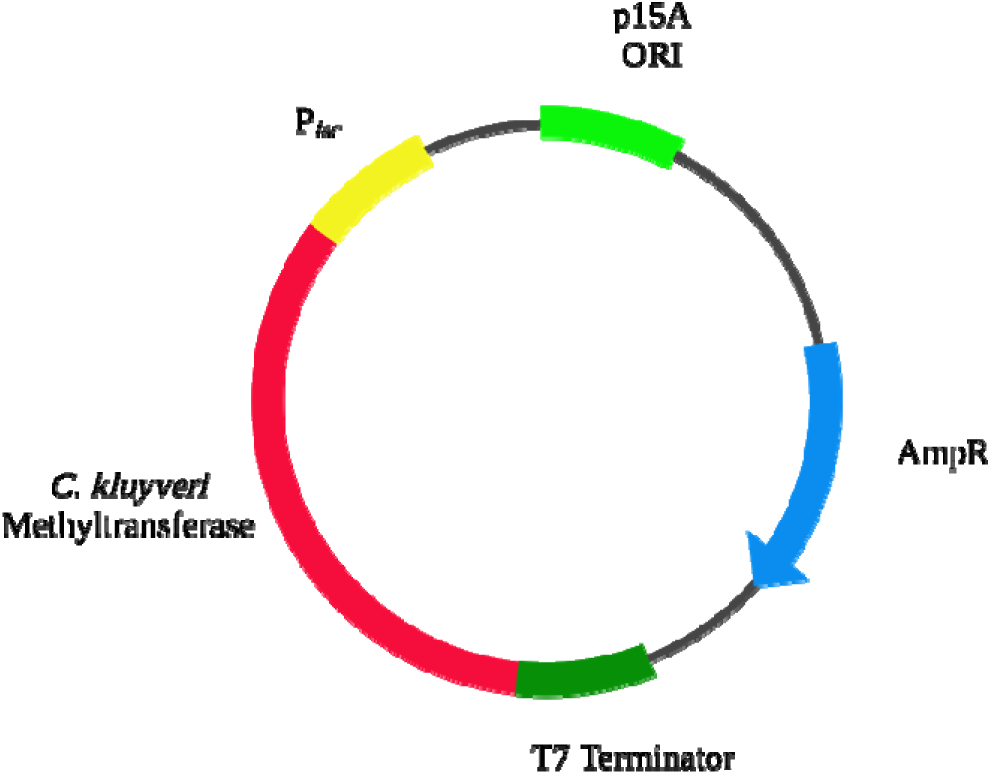
Plasmid map of pMeth based on pUC19. CKL_2671 was amplified from C. kluyveri genomic DNA and cloned into the multiple cloning site of pUC19. The T7 terminator was cloned into the plasmid and the ORI of puC19 was exchanged to the p15A ORI. Created with BioRender.com.

### Triparental conjugational DNA transfer

The *E. coli* TOP10 strain, harboring both pMTL83151 and pMeth plasmids, exhibited reduced growth rates. This growth inhibition was likely attributable to the metabolic burden that was imposed by maintaining resistance to two antibiotics simultaneously (chloramphenicol for pMTL83151 and ampicillin for pMeth), combined with the additional cellular stress of protein expression that is induced by IPTG treatment. To avoid additional burden on this strain by introducing a third plasmid to enable conjugation, we instead performed triparental mating for shuttle vector transfer into the *C. kluyveri* recipient using a modified transformation protocol (38). We retained *E. coli* TOP10, containing both pMeth and the shuttle vector plasmid, as the donor strain and *E. coli* HB101, carrying pRK2013, as the helper strain (39). To support the growth of the *E. coli* strains and *C. kluyveri* on the same plate, we tested the growth of the two *E. coli* strains on RCM plates supplemented with varying ethanol concentrations (86-342 mM, 0.5-2.0% [v/v]) under a N_2_/CO_2_ atmosphere. Both *E. coli* strains exhibited normal growth up to 257 mM ethanol. We decided to use the highest ethanol concentration because it is the closest to the ethanol level that we use to cultivate *C. kluyveri* (342 mM), making it the optimum condition for co-cultivating these three strains.

We evaluated different mating durations (24, 48, 72, 96, and 120 h) by selecting potentially transformed cells on plates containing thiamphenicol and trimethoprim. After 24 h, individual colonies were visible (**Figure S2A**). However, when extending mating times from 48 h to 120 h, individual colonies became indistinguishable on selective plates, requiring re-streaking to isolate single colonies (**Figure S2B)**. In the next step, we successfully verified the presence of the shuttle vector plasmid inside *C. kluyveri* cells *via* colony PCR (**Figure 3A**), and we sequenced the 16S rRNA gene to verify pure cultures (**Figure 3B**). From these findings, we conclude that longer mating times between 48 and 120 h enhance or at least do not adversely affect the transformation efficiency of the cells.

**Figure 3.**
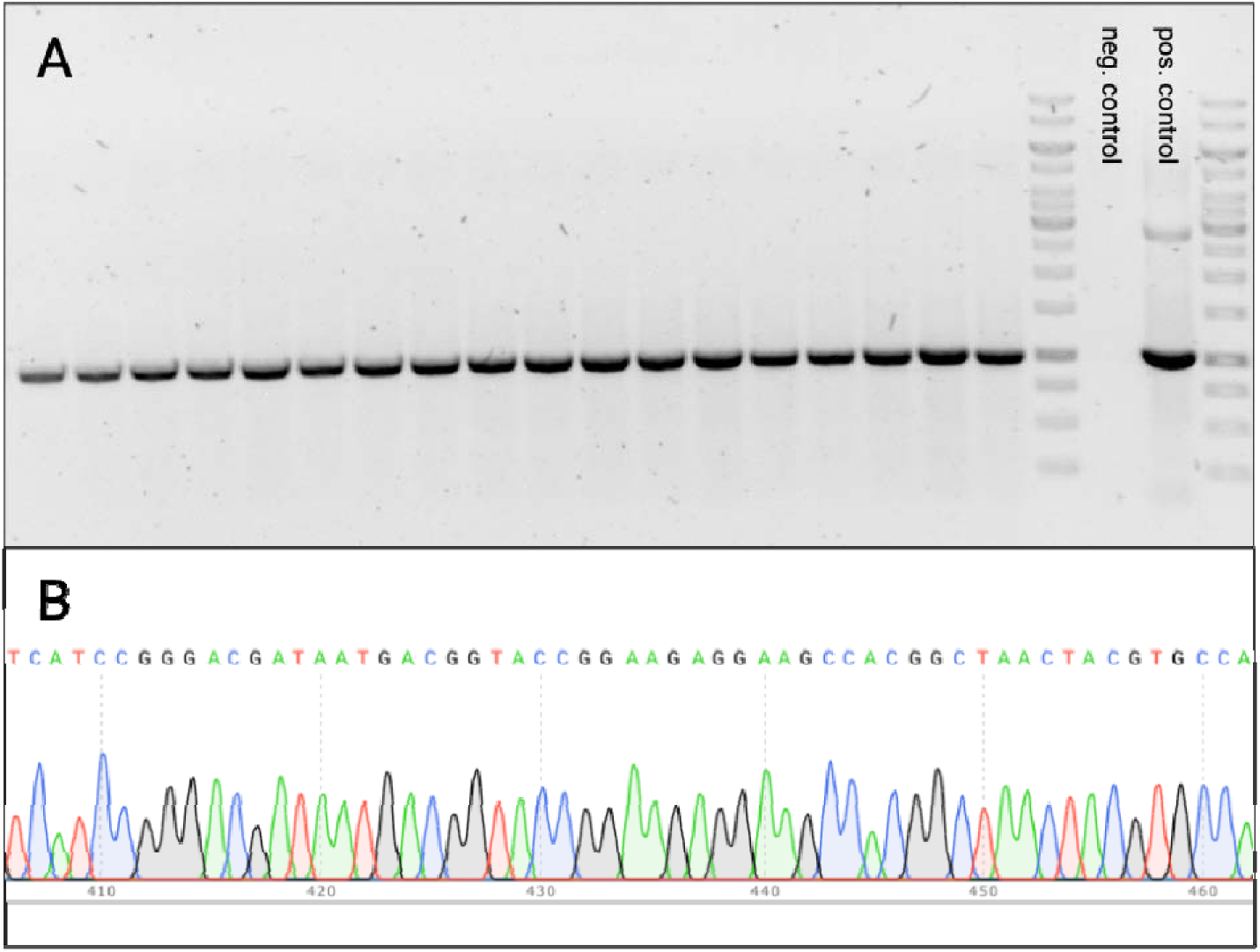
Verification of pMTL83151 inside C. kluyveri cells. A) We screened 18 C. kluyveri colonies (lanes 1-18) for the presence of the plasmid pMTL83151 and verified its presence via colony PCR of the respective clones. Negative and positive (fragment size: ∼1 kbp) controls are shown on the right side. The band corresponding to a longer fragment in the positive control results from the circular pMTL83151 plasmid, which served as template. B) Sanger Sequencing results of the 16S rRNA gene from the screened clones. All clones showed clear signals, confirming pure C. kluyveri pMTL83151 cultures. We show only one of the sequencing results here, which represents all tested clones. Created with BioRender.com.

### Heterologous gene expression as proof of concept

To expand the product portfolio of *C. kluyveri*, we aimed to enable the conversion of *n*-butyrate and *n*-caproate into *n*-butanol and *n*-hexanol. This was achieved by introducing the *adhE2* gene from *C. acetobutylicum* into the shuttle vector plasmid pMTL83151 under the control of its native promoter, which is P*_adhE2_*. *adhE2* encodes a bifunctional alcohol dehydrogenase, which catalyzes both the conversion of acyl-CoA to the corresponding aldehyde in the first step and the reduction of the aldehyde to the respective alcohol in the second step (40). After successful transformation of the plasmid into *C. kluyveri* cells, we observed the production of alcohols corresponding to the chain length of the carboxylates produced. However, preliminary measurements of the alcohols produced by *C. kluyveri* pPadhE2_adhE2 revealed relatively low alcohol concentrations (*n*-butanol: 1.87 mM, *n*-hexanol: 3.15 mM). To increase alcohol production, we tested two conditions: **1)** increasing the expression of the *adhE2* gene by using a stronger promoter (41); and **2)** introducing an NADPH-dependent alcohol dehydrogenase because NADPH represents a more powerful co-factor than NADH (42) and the use of NADH by AdhE2 potentially lowers the NADH/NAD+ ratio too much. Therefore, using NADPH as an alternative co-factor could be advantageous for *n*-butanol and *n*-hexanol production

To address these limitations, we enhanced *adhE2* expression by replacing the native promoter with the reportedly stronger P*_thl_* promoter from *C. acetobutylicum* (43, 44), which resulted in higher *n*-butanol and *n*-hexanol concentrations (*n*-butanol: 2.73 mM, *n*-hexanol: 3.25 mM) compared to the P*_adhE2_* promoter. To further increase alcohol production, we introduced the *bdhB* gene from *C. acetobutylicum*, encoding butanol dehydrogenase B. Unlike AdhE2, BdhB uses NADPH as a co-factor for converting aldehydes to alcohols (45, 46). Since BdhB exclusively catalyzes the second step from the aldehyde to the alcohol (47), we retained AdhE2 to ensure acyl-CoA conversion to the corresponding aldehyde. Thus, we tested three constructs on the shuttle vector plasmid pMTL83151: **1)** pPadhE2_adhE2; **2)** pPthl_adhE2; and **3)** pPthl_adhE2_bdhB and used the pMTL83151 vector as the negative control in a growth experiment (**Figure 4**). We successfully transformed *C. kluyveri* cells with the three constructs and the negative control and evaluated the resulting phenotypes. Notably, during the conjugation process, we observed that DNA transfer for the plasmid encoding both *adhE2* and *bdhB* was only successful after 72 h of mating.

**Figure 4.**
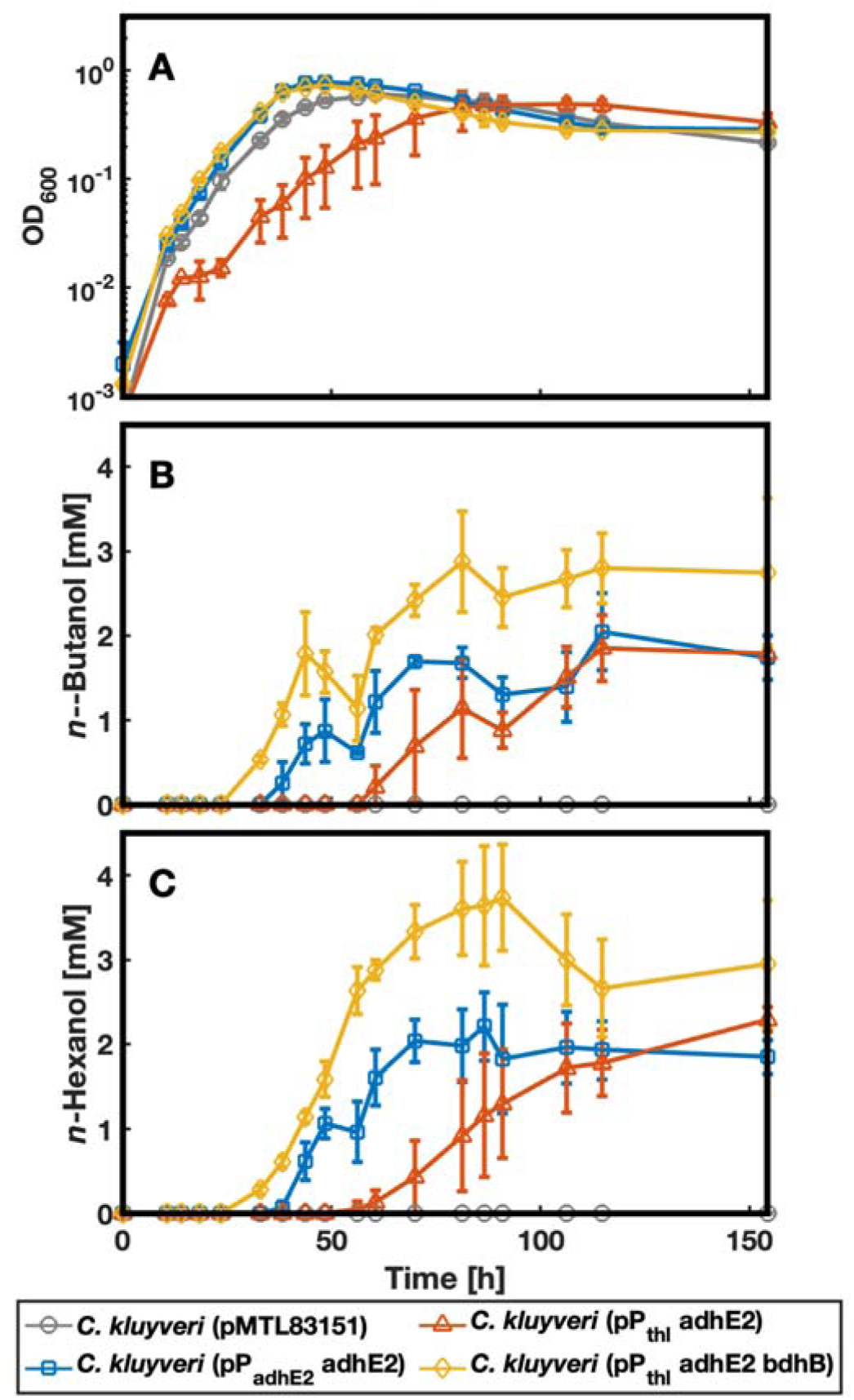
Results of the growth experiment. A) The growth behavior was measured by OD_600_, showing that all strains but C. kluyveri pPthl_adhE2 showed comparable growth. B) C. kluyveri pPthl_adhE2_bdhB reached the highest n-butanol concentration, whereas C. kluyveri pPadhE2_adhE2 and C. kluyveri pPthl_adhE2 showed comparable production levels. C) We observed the highest n-hexanol levels for C kluyveri pPthl_adhE2_bdhB, followed by C. kluyveri pPthl_adhE2. Created with MATLAB2024b.

### *n*-Butanol and *n*-hexanol production of the different strains

In bottle experiments, *C. kluyveri* pPadhE2_adhE2, *C. kluyveri* pPthl_adhE2_bdhB, and the negative control exhibited comparable growth with doubling times between 4.5 and 6 h (**Figure 4A**). In comparison, the three replicates of *C. kluyveri* pPthl_adhE2 started to grow at different times after inoculation, thus, leading to higher standard deviations, not only for the OD_600_ but also for substrate and product concentrations for this strain. However, the growth rates were similar to the other strains tested (data not shown). Unfortunately, several attempts to repeat the growth experiment failed due to a high mutation rate in the promoter region of *adhE2* in *C. kluyveri* pPthl_adhE2. Average *n*-butanol and *n*-hexanol production was first observed after 33 h for *C. kluyveri* pPthl_adhE2_bdhB and after 38 h for *C. kluyveri* pPadhE2_adhE2 (**Figures 4B,C**). Due to the extended lag phase and high standard deviations (up to 54% for *n*-butanol and 173% for *n*-hexanol), *C. kluyveri* pPthl_adhE2 was the last strain starting to produce *n*-butanol and *n*-hexanol (first production observed between 61 h and 81 h, data not shown), resulting in a slower average alcohol production than the other replicates.

Regarding *n*-butanol production, *C. kluyveri* pPthl_adhE2_bdhB reached a maximum concentration of 4.2 ± 0.0 mM, whereas *C. kluyveri* pPadhE2_adhE2 and *C. kluyveri* pPthl_adhE2 exhibited similar *n*-butanol production levels (**Figure 4B**). In line with this observation, *C. kluyveri* pPthl_adhE2_bdhB also achieved the highest *n*-hexanol concentration (3.7 ± 0.6 mM), followed by *C. kluyveri* pPthl_adhE2 (2.3 ± 0.4 mM). *C. kluyveri* pPadhE2_adhE2 produced the lowest amount of *n*-hexanol (2.2 ± 0.4 mM) (**Figure 4C**). The difference in *n*-hexanol production between *C. kluyveri* pPthl_adhE2 and *C. kluyveri* pPadhE2_adhE2 likely would have been notably higher if *C. kluyveri* pPthl_adhE2 had exhibited more consistent growth across the replicates. Also, a longer experiment duration might have further highlighted a potential disparity.

Interestingly, both strains carrying P*_thl_* started producing the corresponding alcohols at lower *n*-butyrate and *n*-caproate concentrations (28.6 ± 10.3 mM and 15.6 ± 12.1 mM, and 34.2 ± 1.4 mM and 21.7 ± 2.3 mM, respectively) (**Figures 4B,C)**. In contrast, *C. kluyveri* pPadhE2_adhE2 started *n*-butanol and *n*-hexanol production at 42.6 ± 2.6 mM *n*-butyrate and 38.7 ± 5.3 mM *n*-caproate (**Figures 4B,C** and **Figures S3C,D)**. Remarkably, *C. kluyveri* pPthl_adhE2_bdhB converted most acyl-CoA into the corresponding alcohol, formed more *n*-butyrate than the other strains, and reached the highest OD_600_ (**Figures 4A** and **Figures S3C,D)**. It produced 30% less *n*-caproate than the other alcohol-producing strains and approximately 35% less than the negative control (**Figure S3D)**. Due to the lower *n*-caproate production, its conversion rate for substrate carbon into product carbon (*n*-butyrate, *n*-caproate, *n*-butanol, and *n*-hexanol) only reached 53%, whereas the other strains converted 62-67% of it (**Figures 4B,C and Figure S3**). Further, *C. kluyveri* pPthl_adhE2_bdhB showed a higher acetate-to-ethanol ratio at the end of the experiment (0.16) than the other strains (0.10-0.12) (**Figures S3A,B)**.

## DISCUSSION

Our results show that: **1)** adapting the growth conditions for *C. kluyveri* on solid DSMZ52 medium; **2)** determining the MIC of thiamphenicol; and **3)** counteracting restriction modification barriers enabled successful DNA transfer of a shuttle vector plasmid into *C. kluyveri* cells *via* conjugation after 24 h of mating. However, extending the mating time to 48 h improved the transformation efficiency. Notably, transferring the plasmid pPthl_adhE2_bdhB (8.234 kbp) was only successful after 72 h, suggesting that shuttle vectors with critical genes or an increased size may require extended mating times. This extension has been shown to improve conjugation efficiencies overall (48).

All generated strains, but the negative control, showed plasmid-based gene expression of the *adhE2* and *bdhB* genes from *C. acetobutylicum*. This led to phenotypically modified *C. kluyveri* strains that were capable of producing *n*-butanol and *n*-hexanol. In line with our expectations, both replacing P*_adhE2_* with the reportedly stronger P*_thl_*promoter and introducing a second, NADPH-dependent butanol dehydrogenase resulted in enhanced *n*-butanol and *n*-hexanol production. The clear variability observed in the lag phase duration among the replicates of *C. kluyveri* pPthl_adhE2 distorted the calculated mean values. This statistical distortion resulted in an artificial lowering of the estimated growth rate because the extended lag phases of some replicates disproportionately influenced the overall growth calculations. This bias not only lowers the slope of the OD_600_ but also needs to be considered for the graphs representing the substrates and products. A potential explanation for the extended lag phase is cell stress caused by the P*_thl_* promoter, which is stronger than P*_adhE2_*. Because AdhE2 catalyzes both reactions of acyl-CoA to aldehyde and aldehyde to alcohol, it may have a higher affinity towards the first reaction in *C. kluyveri*. Hence, the promoter exchange could lead to aldehyde accumulation, which is known to be toxic (49, 50), causing cell stress and impaired cells.

The batch experiment provided compelling evidence supporting the hypothesis that the combination of the P*_thl_* promoter and solely the *adhE2* gene exhibits toxicity. This toxicity was demonstrated conclusively by metabolic impairment: across three independent growth experiments, none of the *C. kluyveri* pPthl_adhE2 replicates produced detectable levels of either *n*-butanol or *n*-hexanol. In contrast, only one experiment resulted in the successful production of the alcohols. Sequencing of the non-alcohol-producing cultures revealed that all contained a deletion in the P*_thl_* region (**Figure S4A**), suggesting a selection pressure against the expression of *adhE2*. Additionally, sequencing *C. kluyveri* pPthl_adhE2 cultures, which produced *n*-butanol and *n*-hexanol, revealed a less defined baseline in the promoter region (**Figure S4B**), indicating that a fraction of the cells had already mutated. Given these findings, we cannot rule out that an unmutated pure culture of *C. kluyveri* pPthl_adhE2 could have performed better. Nonetheless, while using a stronger promoter enhanced the production of *n*-hexanol, adding the butanol dehydrogenase presumably helped accelerate aldehyde removal. This prevented toxic accumulation and resulted in a stable, mutation-free strain.

Furthermore, we observed reduced formation of *n*-caproate by *C. kluyveri* pPthl_adhE2_bdhB despite reaching the highest OD_600_ of all strains. This might be due to a more balanced use of the co-factors NADH and NADPH. Both co-factors are utilized to prevent excessive depletion of NADH, which would result in a reduced NADH/NAD^+^ ratio. This results in a higher NADH reducing capability and increased alcohol production (51). Additionally, improved recycling of the co-factors could explain the lower final concentration of *n*-caproate. Since NADH and NADPH are used as electron donors for alcohol production, they are less available for other processes, such as the formation of carboxylates (28). As a result, these reactions occur less frequently, leading to a decrease in *n*-caproate production.

The genetic system demonstrated in this study enabled plasmid-based gene expression, leading to heterologous production of *n*-butanol and *n*-hexanol by *C. kluyveri*. Besides expanding the product portfolio of this microbe, our research represents the first step towards developing a broader toolkit for *C. kluyveri*, including genome editing techniques (*e.g.*, homologous recombination, CRISPR-based methods). Consequently, the genetic accessibility of the model microbe for chain elongation is a precondition for further research on the reverse ß-oxidation pathway, which holds strong industrial potential. Therefore, gaining a deeper understanding of this pathway, both on a genetic level and in terms of the enzymes involved in their central energy metabolism, is crucial to enhancing its productivity.

## MATERIALS AND METHODS

### Microbial strains, media, and cultivation

*C. kluyveri* DSM555 was obtained from the DSMZ (Braunschweig, Germany) and used as the wild-type strain. Unless stated otherwise, all *C. kluyveri* strains in this study were cultivated in DSMZ52 medium for which 2.5 g/L NaHCO_3_ was used to buffer the medium instead of Na_2_CO_3_. Cultures were grown in 100-mL medium with a headspace of 140 mL under an N_2_/CO_2_ (80/20) atmosphere. Both liquid cultures and plates (containing 0.8% [v/v] Bacto agar) were incubated at 37°C until visible growth was observed. When required, the medium was supplemented with 5 µg/mL thiamphenicol and 10 µg/mL trimethoprim for selection. All genetic manipulations of *C. kluyveri* were conducted under strict anoxic conditions.

Unless otherwise noted, *E. coli* HB101 pRK2013 and *E. coli* TOP10 strains were grown in liquid Luria-Bertani (LB) medium or on LB plates containing 1.5% agar. For plasmid selection and maintenance, the medium was supplemented with 100 µg/mL ampicillin (for liquid cultures), 100 µg/mL carbenicillin (for plates), 30 µg/mL chloramphenicol, or 30 µg/mL kanamycin. IPTG was added at a concentration of 500 mM to induce the expression of the methyltransferase encoded on pMeth.

### Optimization of the plating efficiency

The variables (**Table S1**) were systematically adjusted one at a time to determine the optimal plating conditions, while all other conditions remained identical. The resulting outcomes were compared to the previously identified best conditions. Both the modified condition and the previously best-performing combination of conditions were tested using the same culture conditions and inoculation volume (4% [v/v]) with simultaneous incubation. The inoculum was serially diluted by increasing the dilution factor by a magnitude of 10, reaching a final dilution of 1:100,000, and each dilution was plated in duplicates for both conditions. Colony counts were performed by capturing images using the UVP GelStudio system (Analytic Jena, Jena, Germany) and analyzed with the integrated colony-count software. The plating conditions yielding the highest colony count were determined to be the updated optimal combination, and the following variable was tested accordingly.

### Restriction digest

10 mL of a late exponential phase *C. kluyveri* culture were harvested, washed twice with phosphate-buffered saline (PBS) buffer (137 mM sodium chloride, 2.7 mM potassium chloride, 10 mM sodium phosphate dibasic, and p1.8 mM potassium phosphate monobasic), and resuspended in 300 µL PBS buffer. Cells were lysed using bead-beating (30 sec, 6.5 m/s; FastPrep-24^TM^, MP BioMedicals, Santa Ana, USA). The lysing procedure was performed four times, and in between, the cells were kept on ice for 3 min. The lysis was centrifuged at 2000 X *g* for 20 min at 4°C. To assess the sensitivity of the respective plasmid to the native restriction enzymes of *C. kluyveri*, 5 µL of the supernatant were added to 13 µL plasmid (200 ng) and 2 µL rCutSmart^TM^ buffer (New England Biolabs, Ipswich, Massachusetts, USA). The prepared reaction mix was stored at 37°C for 1 to 24 h. Results were analyzed *via* gel electrophoresis and potential corresponding restriction sites were analyzed *in silico* with SnapGene (GSL Biotech LLC, San Diego, USA).

### PacBio sequencing

High molecular weight DNA (25 ng/µL) was isolated with the NucleoSpin Microbial DNA Mini kit for DNA (Macherey-Nagel, Düren, Germany) and sheared using 35 kb settings with a Megaruptor 2 instrument (Diagenode, SA, Lüttich, Belgium). Average insert sizes were approximately 10 kb. Sheared fragments were used to prepare libraries with the HiFi SMRTbell Express Template Prep Kit 2.0 (Pacific Biosciences, Menlo Park, California, USA). The libraries were size-selected with a BluePippin system (SageScience, Beverly, USA) with 17 kb cutoff in a 0.75% DF Marker S1 High-Pass 6-10 kb vs3 gel cassette (Biozym, Hamburg, Germany). The library was sequenced with sequencing primer v2 (Pacific Biosciences, #101-847-900) and 4 h of pre-extension time on a single SMRT Cell with the Sequel II system using Binding Kit 2.0.

### Methylation profiling

Reads were demultiplexed using Lima (52) and converted into High-Fidelity (HiFi) reads retaining kinetic information with Circular Consensus Sequencing (53). BAM files were converted to FASTQ with bam2fastx (54). Reads shorter than 1000 bp and the lowest 5% of reads based on quality scores were discarded using Filtlong (55). Trycycler (56) was used to subsample reads and generate a consensus sequence from 12 assemblies independently generated with Flye (57), Miniasm (58), and Raven (59). HiFi reads were then aligned with the consensus assembly using pbmm2 (60), and ipdSummary (61) was used to detect and classify DNA base-modifications from the kinetic signatures. MotifMaker (62) was used to identify statistically significant methylated motifs, with a minimum score threshold of 60. Motifs were retained if they were identified for 90% or more possible sites in the genome. The native methyltransferase of *C. kluyveri,* responsible for methylating the critical CCGG sites, was identified using the REBASE database (37).

### Plasmid construction

Plasmids (**Table S2**) were constructed with *E. coli* TOP10 and the respective primers (**Table S3)**. The pUC19 vector (63) served as the backbone for the methylation plasmid pMeth. The gene encoding the critical methyltransferase (CKL_2671) from *C. kluyveri* was PCR amplified (Q5 polymerase, New England Biolabs (NEB), Ipswich, MA, USA) from genomic DNA (NucleoSpin, Microbial DNA, Macherey-Nagel) and inserted into the multiple cloning site *via* Gibson Assembly (NEBuilder, NEB). The lambda t1 terminator was introduced downstream of the CKL_2671 gene *via* PCR, amplifying the plasmid with overlapping primers and transferring the linear DNA inside *E. coli*. The same method was applied to remove the nucleotide sequence between the promoter and the start codon of CKL_2671 to utilize the promoter present in the backbone (P*_lac_*). In the final step, the origin of replication (ORI) was exchanged from the pSC101 ORI to the p15a ORI (64) by Gibson Assembly (NEBuilder, NEB) to ensure a low copy number and compatibility with the shuttle vector plasmids for *C. kluyveri*. For *n*-butanol and *n*-hexanol production, pMTL83151 (35) served as the backbone for all shuttle vector plasmids. The DNA fragments *adhE2*, *bdhB*, P*_adhE2_*, and P*_thl_* from *C. acetobutylicum* were PCR amplified (Q5 polymerase, NEB) from genomic DNA, and the respective fragments were inserted into the plasmids *via* Gibson Assembly (NEBuilder, NEB).

### Triparental conjugational DNA transfer

We inoculated LB medium supplemented with ampicillin, chloramphenicol, and IPTG (to induce the expression of the methyltransferase) with the donor strain *E. coli* TOP10, carrying the desired plasmid and incubated the culture overnight. When both the donor strain and the helper strain *E. coli* HB101 pRK2013, which was grown in LB medium supplemented with 30 µL kanamycin, reached an OD_600_ of 0.6-0.8, 1 mL of each culture was harvested by centrifugation (17,900 X *g*, 3 min). The resulting pellets were suspended and mixed in PBS buffer. We again centrifuged the *E. coli* mixture (17,900 X *g*, 3 min) and transferred the received cell pellet into the anoxic chamber. Subsequently, 2 mL of mid-exponential *C. kluyveri* wild-type culture (OD_600_: 0.4-0.7) was harvested *via* centrifugation (9.800 X *g*, 4 min) and resuspended in 50 µL RCM under an N_2_ atmosphere. This cell suspension was used to resuspend the *E. coli* cell pellet. The cell mixture was then spot-plated on RCM plates (1.5% [v/v] agar) containing 500 mM IPTG and 257 mM ethanol (1.5% [v/v]). Plates were incubated at 37°C in an N_2_/CO_2_ (80/20) atmosphere for up to 120 h. After mating, cells from the mating spots were transferred *via* striking onto DSMZ52 plates containing thiamphenicol to select for the respective plasmid. After cells had grown on the first selective plate, they were streaked onto a thiamphenicol and trimethoprim-containing DSMZ52 plate to select against *E. coli* strains. The resulting colonies were then further analyzed.

### Molecular methods for analysis of genetically modified *C. kluyveri*

Direct colony PCR was conducted on both colonies and liquid cultures to confirm the presence of the respective plasmid in *C. kluyveri* cells. Individual colonies were picked and resuspended in 25 µL of 10 mM NaOH to extract the DNA template. For PCR performed on liquid cultures, 100 µL of culture was sampled. In both cases, the cells were lysed at 98°C for 10 min using a ThermoMixerC (Eppendorf, Hamburg, Germany). For the PCR reactions, 1 µL of the lysate was utilized in two separate 10 µL PCR reactions (PhirePlant (ThermoScientific^TM^, Waltham, USA)) – one using plasmid-specific primers and the other using *C. kluyveri*-specific primers. The results of the reactions were analyzed *via* gel electrophoresis. Additionally, a 30 µL PCR reaction targeting the 16S rRNA gene was performed using 3 µL of the lysate as a template. The resulting PCR product was purified using the QIAquick PCR Purification Kit (QIAGEN, Venlo, Netherlands) and subsequently analyzed *via* Sanger sequencing (GENEWIZ, Leipzig, Germany) to verify a pure culture of *C. kluyveri*.

### Analysis of compound concentration

To determine the concentrations of carboxylates (*e.g.*, acetate, *n*-butyrate, *n*-caproate) and alcohols (*n*-butanol and *n*-hexanol), an Agilent 7890B gas chromatograph (Agilent Technologies, Inc., Santa Clara, CA, USA) that were equipped with a DB-Fatwax UI capillary column (30 m x 0.25 mm x 0.25 µm) and flame ionization detector (FID) was used. Hydrogen served as the carrier gas, and the temperature program consisted of an initial temperature of 40°C for 3 min, ramped at 12°C/min to 160°C, followed by a further ramp of 20°C/min to 230°C with a 5-min hold. Ethyllactate was used as an internal standard at a concentration of 2 mM for quantifying carboxylates. The sampling volume was 1 mL of culture that was centrifuged (17,900 *g*, 3 min), and from which we stored the supernatant at −20°C until further analysis. Before measuring, the samples were thawed and filtered through a 0.2 µm polyvinylidene difluoride (PVDF) sterilefilter. For the analysis, samples were diluted at ratios of 1:2 for *n*-butanol and *n*-hexanol and 1:50 for MCCs measurements.

For ethanol concentration measurements, undiluted samples were analyzed using the LC-20 high-performance liquid chromatography (HPLC) system (Shimadzu, Nishinokyo Kuwabara-cho, Japan). The system was equipped with an Aminex-HPX-87H column, and 5 mM sulfuric acid was utilized as the mobile phase at a flow rate of 0.6 mL/min (LC-20AD). The oven temperature was maintained at 65°C (CTO-20AC), and the samples were cooled at 15°C in the autosampler unit.

## ACKNOWLEDGEMENTS

We want to express our sincere gratitude to Silke Dauser and Heike Budde for performing the library preparation and PacBio sequencing. Further, the authors want to thank José Antonio Velázquez Gómez for supporting the HPLC analysis. Funding sources were the Max Planck Institute to L.T.A. as a Max Planck Fellow, the Alexander von Humboldt Foundation through the Alexander von Humboldt Professorship (L.T.A.), the Novo Nordisk Foundation through the CO_2_ Research Center (CORC) with grant number NNF21SA0072700 (L.T.A.), the Deutsche Forschungsgesellschaft (DFG, German Research Foundation) through the Leibniz Prize (L.T.A.), and the Deutsche Forschungsgemeinschaft under Germany’s Excellence Strategy – EXC 2124 – 390838134 (L.T.A.).

L.T.A. initiated the work. C.S., S.T.B., and L.T.A. designed the experiments. C.S., S.T.B., J.W.M., and K.G. performed laboratory experiments and analyzed the data. L.T.A. and S.T.B. supervised the project. C.S., L.T.A., J.W.M., and S.T.B. wrote the manuscript, while all edited the paper and approved the final version. We declare no conflict of interest.

## SUPPORTING INFORMATION

### Plating efficiency

The plating efficiency was optimized by stepwise exchanging one condition while using the same pre-culture for each comparison (**Table S1**). Colony forming units (CFU) were counted, and the best-performing combination of the two combinations was defined as the current best condition. The current best condition was then used as a control for the next round of the experiment when, again, one factor/condition was changed. If the following combination showed better results than the control, it was updated to be the new current best condition. As a result, we achieved an iterative improvement of the plating efficiency in each round. We observed the biggest positive impact on the plating efficiency by supplementing a 10x concentration of the seven-vitamin solution, compared to the concentration used for cultivating *C. kluyveri* in liquid culture. We refer to this modified medium as DSMZ52*. The different conditions that we tested and their respective outcome are shown in **Table S1**.

**Table S1.** Optimization of the plating efficiency. Each line represents an independent and individual experiment in which two conditions were tested, respectively. Experiments are listed chronologically, meaning that the plating efficiency stays constant with every line or is improved. DSMZ52* means that DSMZ52 medium with 10x vitamin supplementation was used. From all conditions tested, pour plating a culture in its early exponential phase in DSMZ52* with 0.8% agar, 0.6 bar overpressure (N_2_/CO_2_), and a filter paper soaked with ethanol showed the best plating result.

| Dilution factor | Current best combination | CFU per plate | Adapted condition | CFU per plate |
| --- | --- | --- | --- | --- |
| 1:100 | Spread plating on DSMZ52 | 563 | Spread plating on DSMZ52 with 10x vitamin supplementation | 1,095 |
| 1:1,000 | Spread plating on DSMZ52* | 707 | Spread plating on DSMZ52*, with paperclips for increased ventilation of CO2 | 148 |
| 1:100 | Spread plating on DSMZ52* | 1,001 | Spread plating on RCM | 402 |
| 1:1,000 | Spread plating on DSMZ52* | 980 | Pour plating in DSMZ52* in 0.8% agar | 1,056 |
| 1:100 | Pour plating in DSMZ52* in 0.8% agar | 944 | Pour plating in DSMZ52* in 0.8% agar, additional EtOH in the headspace | 141 |
| 1:10,000 | Pour plating in DSMZ52* 0.8% agar, early exponential phase | 678 | Pour plating in DSMZ52* in 0.8% agar, late exponential phase | 216 |
| 1:10,000 | Pour plating in DSMZ52* in 0.8% agar, early exponential phase | 577 | Pour plating in DSMZ52* in 0.8% agar, early exponential phase, at 0.6 bar overpressure | 748 |
| 1:10,00 | Pour plating in DSMZ52* in 0.8% agar, early exponential phase, at 0.6 bar overpressure | 551 | Pour plating in DSMZ52* in 0.8% agar, early exponential phase, at 0.6 bar overpressure, with a filter paper soaked with EtOH inside the petri dish | 579 |

\*CFU: colony forming units per plate
\*DSMZ52\*: DSMZ52 medium with 10x seven-vitamin solution

### Determination of the minimal inhibitory concentration of thiamphenicol for *C. kluyveri*

Wild-type cells of *C. kluyveri* were grown in liquid DSMZ52 medium at different thiamphenicol concentrations to determine the minimum inhibitory concentration (MIC). After one week, we observed growth up to a thiamphenicol concentration of 2 µg/mL, suggesting that 3 µg/mL was the MIC (**Figure S1**). However, to ensure no wild-type cells would grow, for example, due to spontaneous resistance, we decided to use a thiamphenicol concentration of 5 µg/mL to select the plasmid-carrying cells in liquid medium and 3 µg/mL on plates.

**Figure S1.**
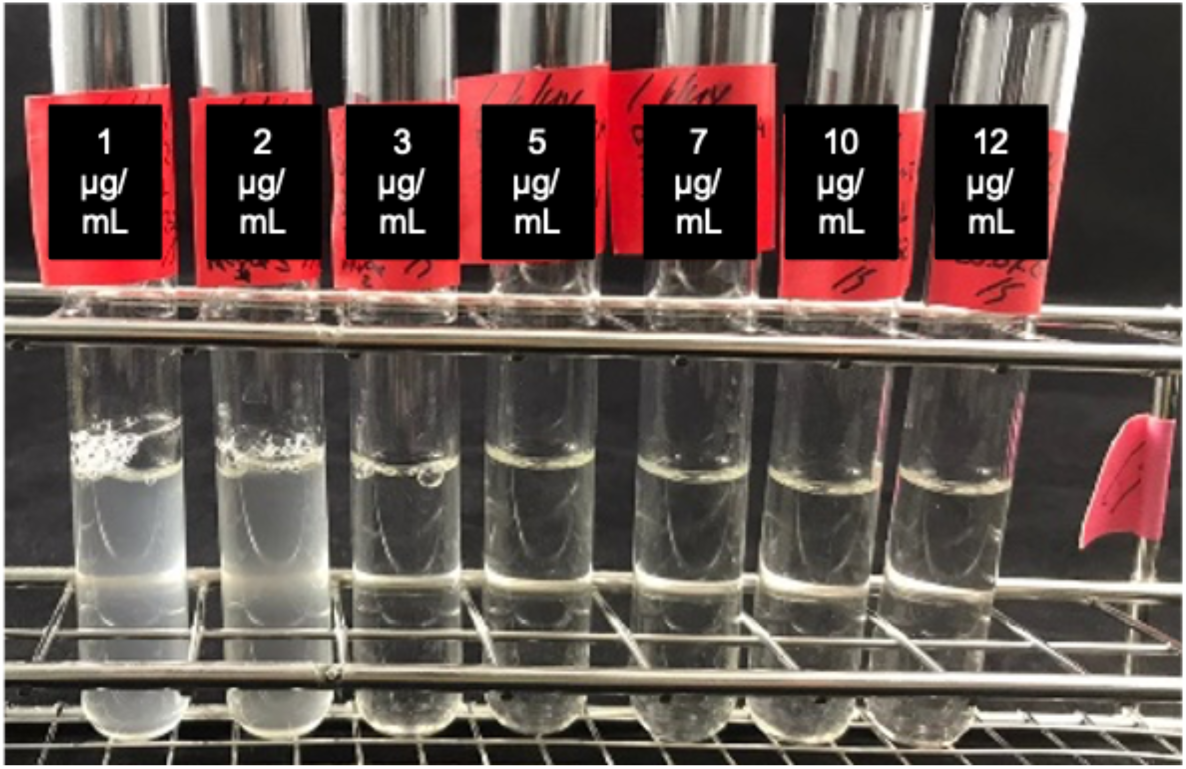
Determination of the MIC of thiamphenicol for C. kluyveri.

### Selection for plasmids

To identify the ideal mating time to obtain *C. kluyveri* mutants, we restreaked the potentially pMTL83151-carrying cells after 24, 48, 72, 96, and 120 h of mating. After 24 h, we observed individual colonies (**Figure S2A**), whereas, after 48 h of mating, the plates showed a lawn of cells carrying the plasmid (**Figure S2B**), indicating that more of the transferred cells carried the plasmid pMTL83151. Because colonies could not be distinguished after 48 h for the empty vector pMTL83151, no increased efficiency could be observed after more than 48 h of mating. However, longer mating times might be beneficial for larger or more toxic plasmids.

**Figure S2.**
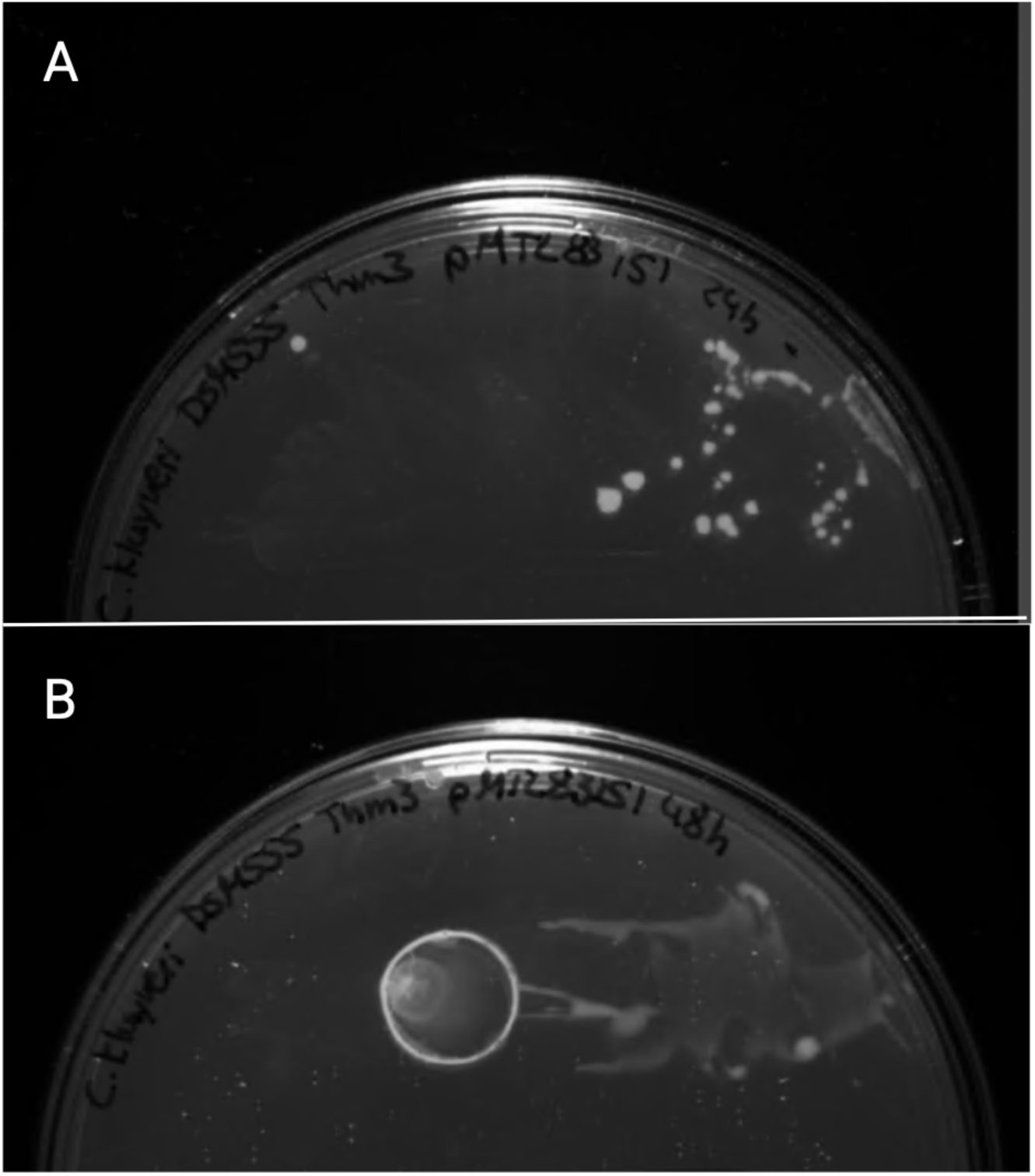
Growth of C. kluyveri colonies on selective plates after 24 and 48 h mating time.

### Growth experiment for *n*-butanol and *n*-hexanol production

We measured the substrates, ethanol (**Figure S3A**) and acetate (**Figure S3B**), and the products, *n*-butyrate (**Figure S3C**) and *n*-caproate (**Figure S3D**), for the different mutant strains (*C. kluyveri* pMTL83151, *C. kluyveri* pPadhE2_adhE2, *C. kluyveri* pPthl_adhE2, and *C. kluyveri* pPthl_adhE2_bdhB producing *n*-butanol and *n*-hexanol and the negative control carrying the empty pMTL83151 vector throughout time.

**Figure S3.**
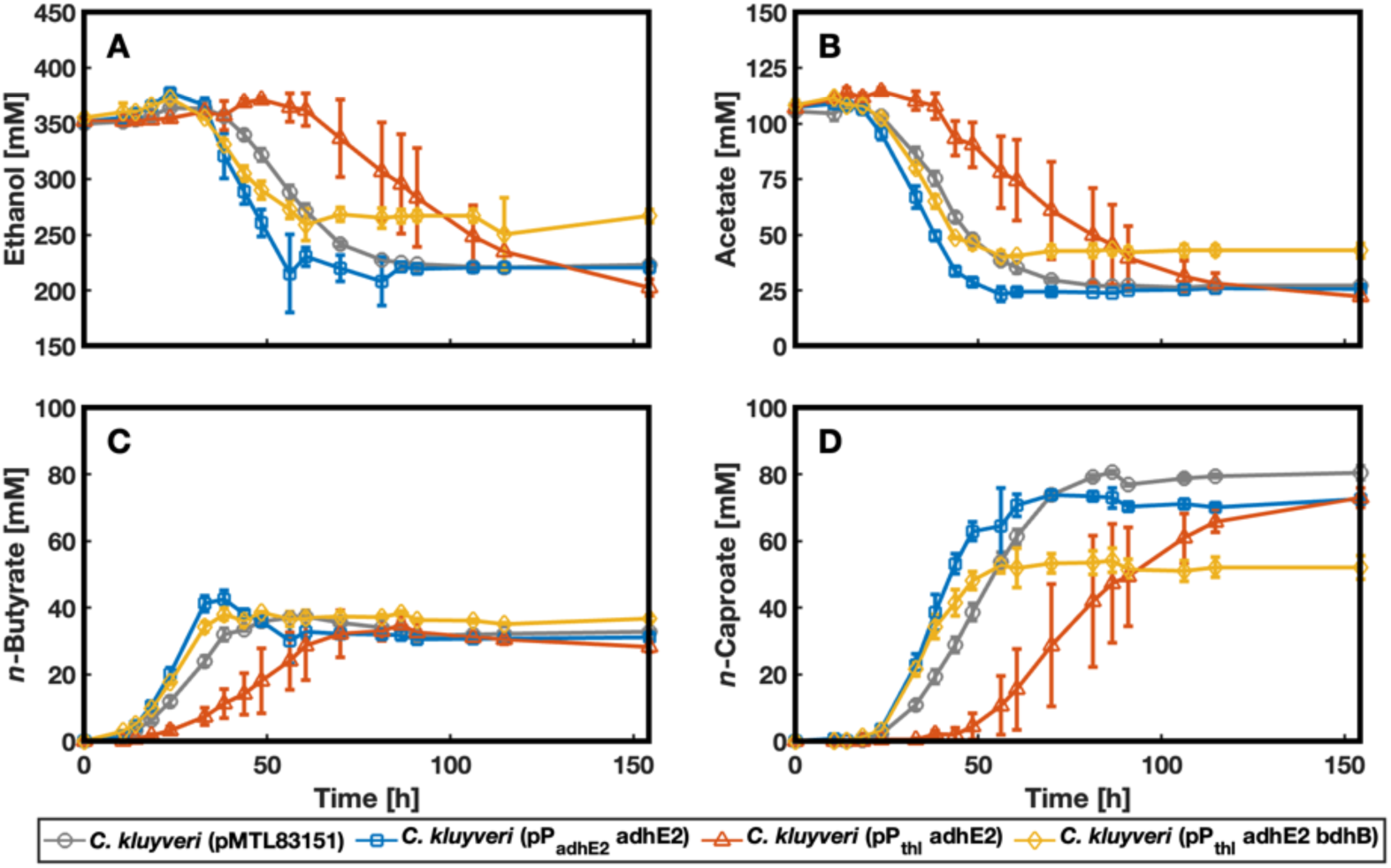
Concentration of the substrates ethanol A) and acetate B) and the products n-butyrate C) and n-caproate D).

### Sanger Sequencing of P*thl* in *C. kluyveri* pPthl_adhE2 cells

Sanger Sequencing of *C. kluyveri* pPthl_adhE2 revealed mutations in the promoter region *P_thl_* of the *C. kluyveri* pPthl_adhE2, which did not produce any longer-chain alcohols than ethanol (**Figure S4A**). Also, the replicates of *C. kluyveri* pPthl_adhE2 seemed to have a fraction of mutated cells (marked with the red box) at the end of the experiment (**Figure S4B**).

**Figure S4.**
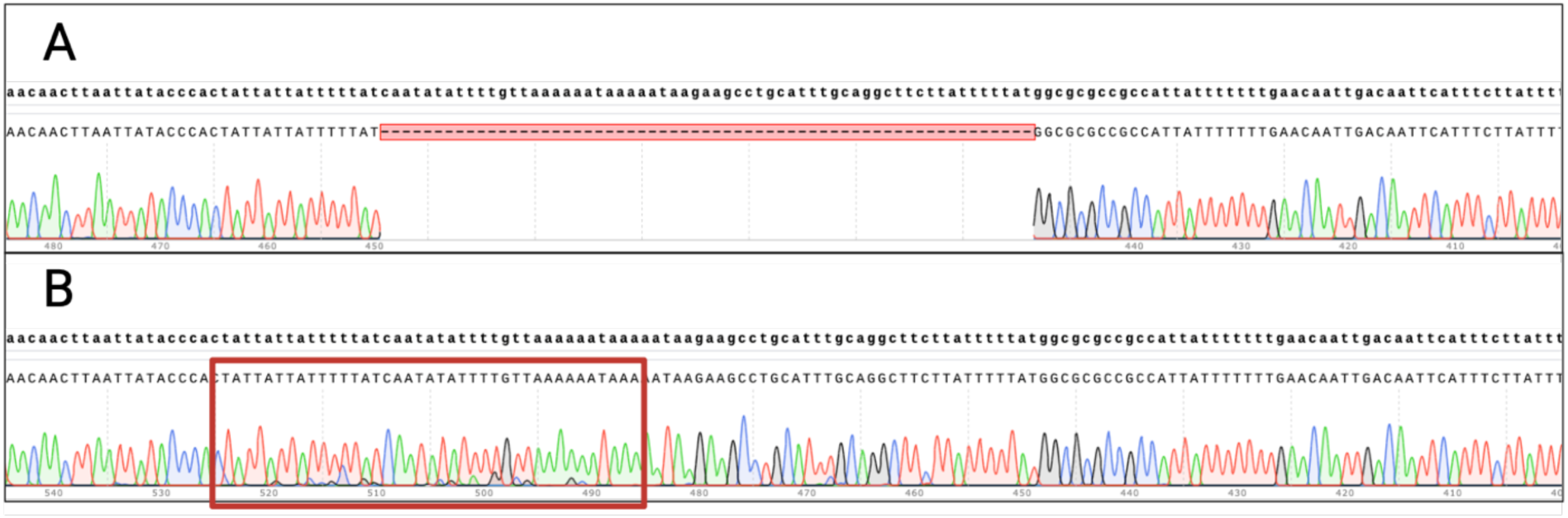
Sequencing results of the promoter region of C. kluyveri pPthl_adhE2. A) Deletion in the promoter region (P_thl_) of adhE2 in C. kluyveri pPthl_adhE2 cultures, which did not produce n-butanol and n-hexanol. B) Shows the same, not deleted, promoter region of C. kluyveri pPthl_adhE2 cultures, which produced n-butanol and n-hexanol, at the end of the experiment. However, sequencing these cultures revealed mixed signals (especially inside the region marked with the red box), indicating that a fraction of the cells had mutated during the growth experiment. Created with BioRender.com.

### Plasmid list

**Table S2.**
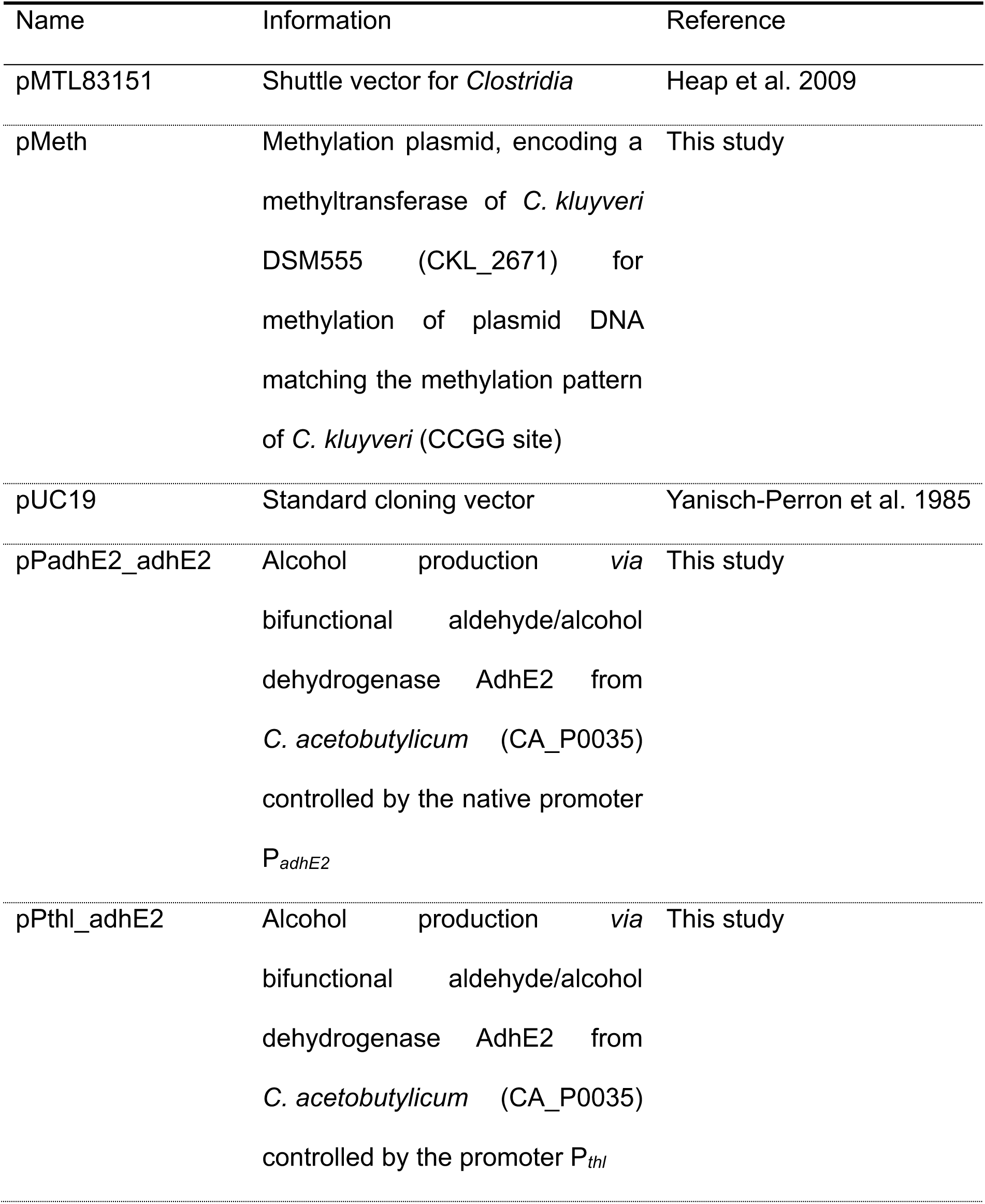
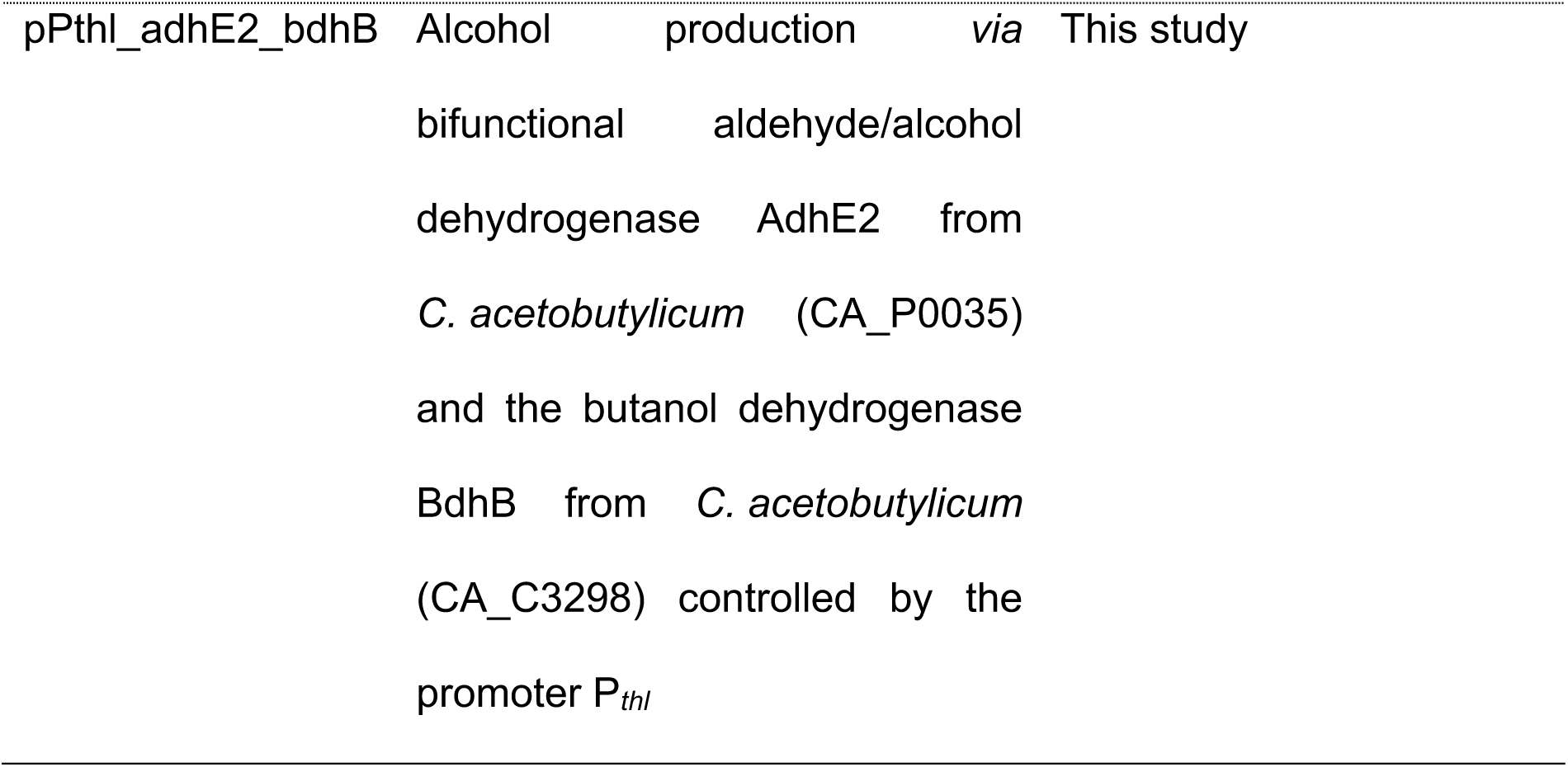
List of plasmids used in this study.

### Primer list

**Table S3.**
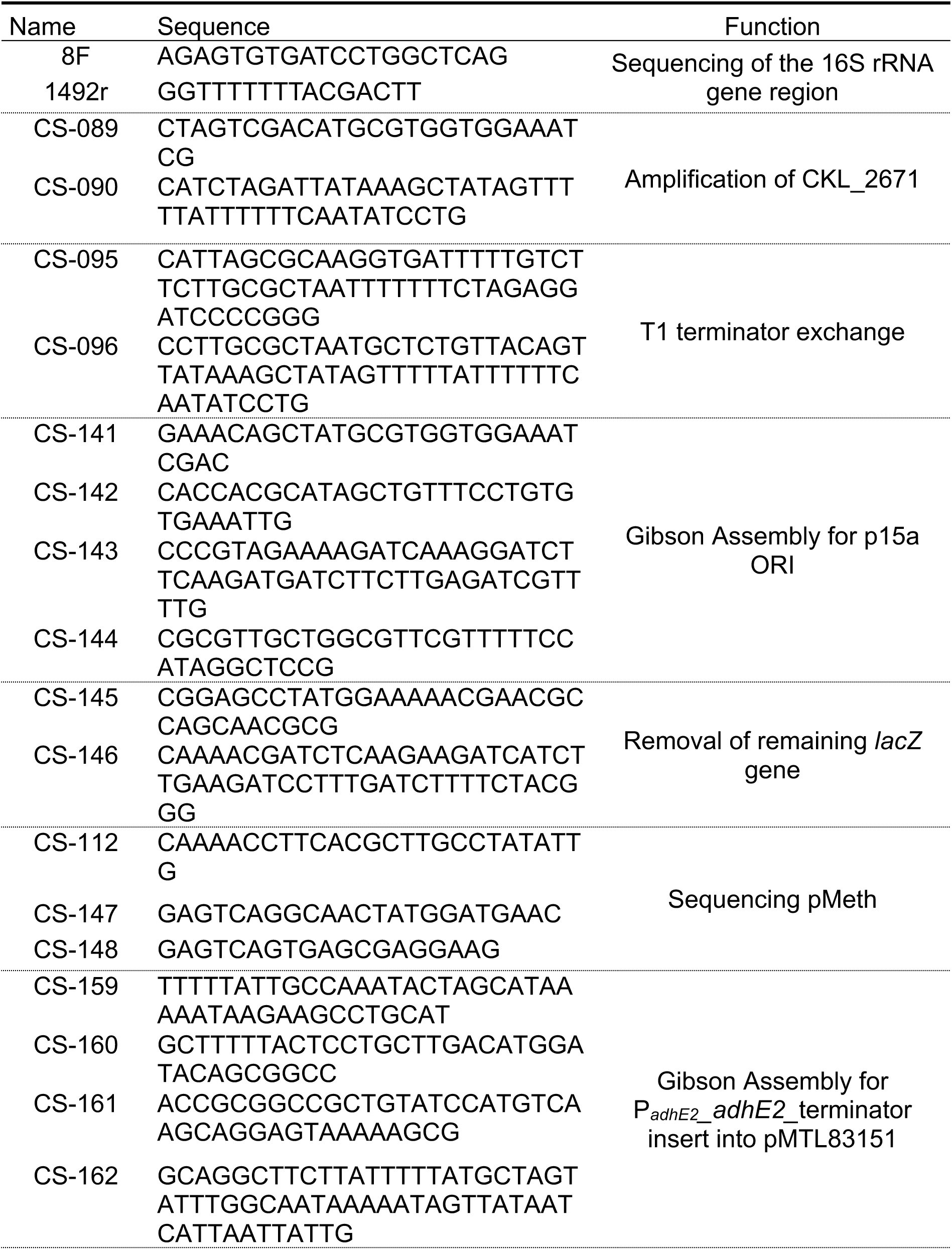
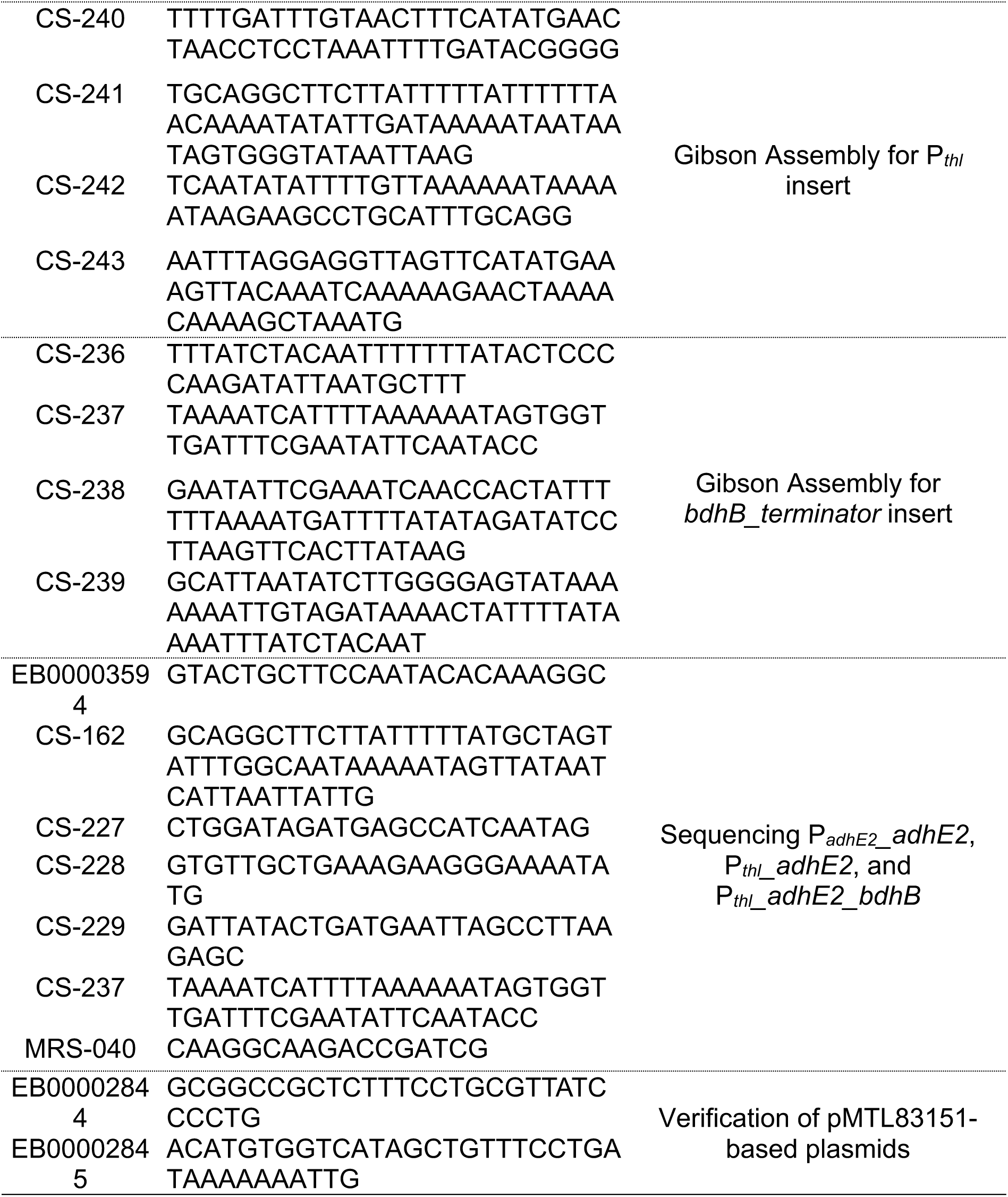
List of oligonucleotides used in this study.

### Bacterial strains

**Table S4.**
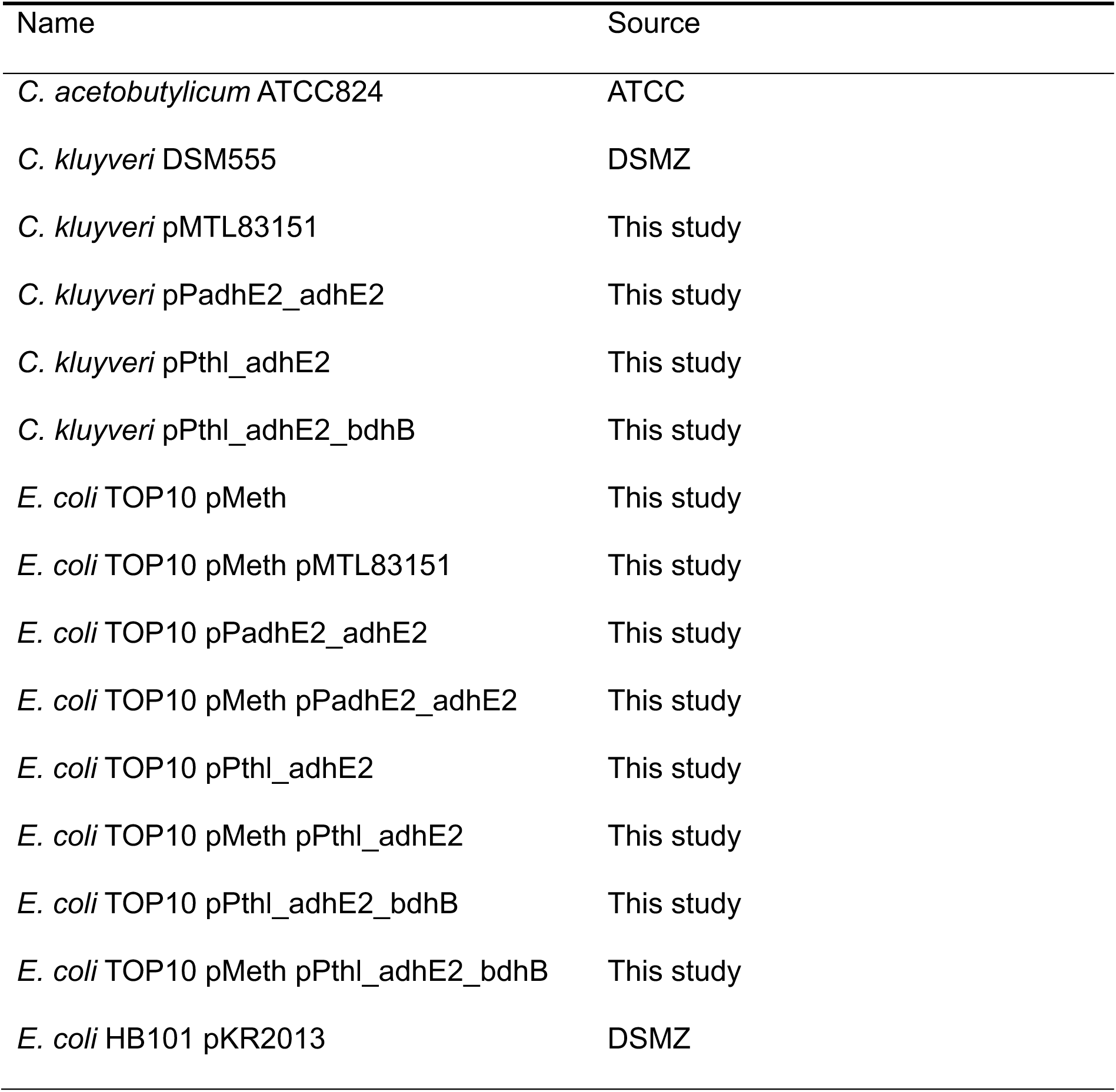
List of bacterial strains used in this study.

